# Age-related morphological changes in the *C57BL/6J* mouse calvaria: implications for stereotaxic neurosurgery

**DOI:** 10.64898/2026.04.24.720718

**Authors:** Zachary A. Knight, Heeun Jang

**Affiliations:** Department of Physiology; University of California San Francisco, San Francisco, CA 94158, USA; Kavli Institute for Fundamental Neuroscience; University of California San Francisco, San Francisco, CA 94158, USA; Howard Hughes Medical Institute; University of California San Francisco, San Francisco, CA 94158, USA; Department of Biomedical Sciences, College of Veterinary Medicine, Iowa State University, Ames, IA 50011, USA

**Keywords:** calvaria, aging, stereotaxic surgery, skull morphometry, sexual dimorphism, C57BL/6J, bregma, lambda

## Abstract

The mouse skull serves as the coordinate framework for stereotaxic neurosurgery, yet whether aging alters surgically critical calvarial geometry has not been examined. We measured three parameters in large cohorts of young (2-3 months) and aged (23-30 months) male and female C57BL/6J mice: bregma-lambda distance and the dorsoventral depth at rostral-lateral (X = ± 2.5 mm, Y = 0) and caudal-lateral (X = ± 2.5 mm, Y = −0.65 mm) calvarial sites. Bregma-lambda distance was significantly reduced with aging in both sexes. Similarly, the rostral-lateral calvaria showed age-related flattening in both male and female mice. The more caudal-lateral site showed smaller but significant age-related flattening, with males exhibiting significantly greater reductions than females. Aging increased caudal-lateral shape variability while reducing bregma-lambda distance variability in males. On the contrary, aging reduced rostral-lateral shape variability in females. These findings demonstrate that the mouse skull undergoes significant, spatially non-uniform, and sexually dimorphic remodeling into old age, where skull shrinks along the midline and flattens more significantly at the rostral than caudal plane. Stereotaxic protocols designed for young adult mice may systematically misestimate skull geometry in aged animals. Age-stratified reference data should be incorporated into stereotaxic practice for aged rodent models.

## INTRODUCTION

The mouse calvaria serves as the coordinate reference system for stereotaxic neurosurgery. Targeting of brain structures depends on the positional stability of two cranial landmarks, bregma and lambda, and on the assumption that lateral calvarial points lie in a predictable dorsoventral (Z) plane (Cecyn and Abrahao 2023; Paxinos and Franklin 2019). While standard atlases are calibrated on young adult (2-4 month) mice, use of aged animals (18-30 months) are now increasingly important to study neurodegeneration and neuroinflammation (Klæstrup et al. 2022; Drechsler et al. 2016; Nazem et al. 2015). Increasing studies use naturally aged rodents for stereotaxic brain surgeries (Tokizane et al. 2024; Li et al. 2022; Tryon et al. 2020; Friedman et al. 2020; Ogbeide-Latario et al. 2022; Jang et al. 2024). However, whether aging systematically alters the skull geometry on which these atlases depend has not been examined.

The craniofacial skeleton undergoes continuous remodeling throughout adulthood driven by shifts in the osteoblast-osteoclast balance, declining sex-steroid signaling, and progressive osteoblast apoptosis (Lee et al. 2026; Corrado et al. 2020; Manolagas 2010). In humans, aging reduces calvarial volume, shortens the sagittal diameter, and produces region-specific changes in vault shape, with morphological effects showing sexual dimorphism (Walczak et al. 2023; Cotofana et al. 2018). In mice, aging involves losses of calvarial neurovascular density (Horenberg et al. 2025), consistent with progressive periosteal remodeling. Differences in skull morphological landmark distances and skull bone thickness that depend on age and sex have been reported within young adult mice (Gulner et al. 2022). However, the macroscopic shape changes directly relevant to surgical landmark geometry, such as bregma-lambda distance and lateral calvarial height, have not been quantified in aged mice.

Inter-animal variability in bregma and lambda position is a recognized source of stereotaxic targeting error (Cecyn and Abrahao 2023; Rangarajan et al. 2016). If aging systematically alters these coordinates, atlas-derived coordinates may be inaccurate in aged cohorts. Here, we quantify age-related changes in three surgically critical calvarial parameters: bregma-lambda distance and the Z coordinates at rostral-lateral and caudal-lateral sites, in large cohorts of male and female C57BL/6J mice, the most commonly used strain in neuroscience research, in 2-3 and 23-30 months of age.

## MATERIALS AND METHODS

### Animals

All procedures were approved by the Institutional Animal Care and Use Committee (IACUC) of University of California, San Francisco and conducted in accordance with the *Guide for the Care and Use of Laboratory Animals* (8th ed., National Academies Press, 2011). Male and female C57BL/6J mice (The Jackson Laboratory, No. 000664) were housed in groups of up to 5 per cage in temperature- and humidity-controlled facilities under a 12-h light/dark cycle with *ad libitum* access to water and standard chow (PicoLab Rodent Diet 20, #5053). Young adult mice were studied at 2-3 months of age; aged mice were housed until 23-30 months of age before measurement.

### Cranial coordinate measurements

Mice were anesthetized with 5% isoflurane in a chamber and maintained at 1-2% isoflurane delivered via nose cone, and secured in a stereotaxic frame (Kopf Instruments, Model 942 and 922) (**Figure 1A**). The scalp was incised along the midline and residual periosteum was gently removed; the exposed skull was cleaned with sterile saline and allowed to dry. Bregma was defined as the point of intersection of the coronal and sagittal sutures, and lambda as the point of intersection of the lambdoid and sagittal sutures, following the conventions (Paxinos and Franklin 2019; Cecyn and Abrahao 2023). When sutural intersections were not clearly defined, bregma and lambda were identified by extrapolating smooth hypothetical lines along the respective sutures. A small pilot hole (∼0.2 mm diameter) was drilled at bregma, which was designated the stereotaxic origin [X = 0, Y = 0, Z = 0]. The bregma-lambda distance was recorded as the anteroposterior (Y-axis) span from bregma to lambda with the arm zeroed in Z at bregma. The dorsoventral (Z) coordinate was then measured at X = ± 2.5 mm, Y = 0 (rostral-lateral site) and X = ± 2.5 mm, Y = - 0.65 mm (caudal-lateral site) by lowering the stereotaxic arm until it contacted the skull surface. All measurements were performed by a single operator to minimize personnel-dependent variability. After completion of measurements, animals underwent survival surgeries for viral injection and/or optic fiber implantation and were allowed to recover.

**Figure 1.**
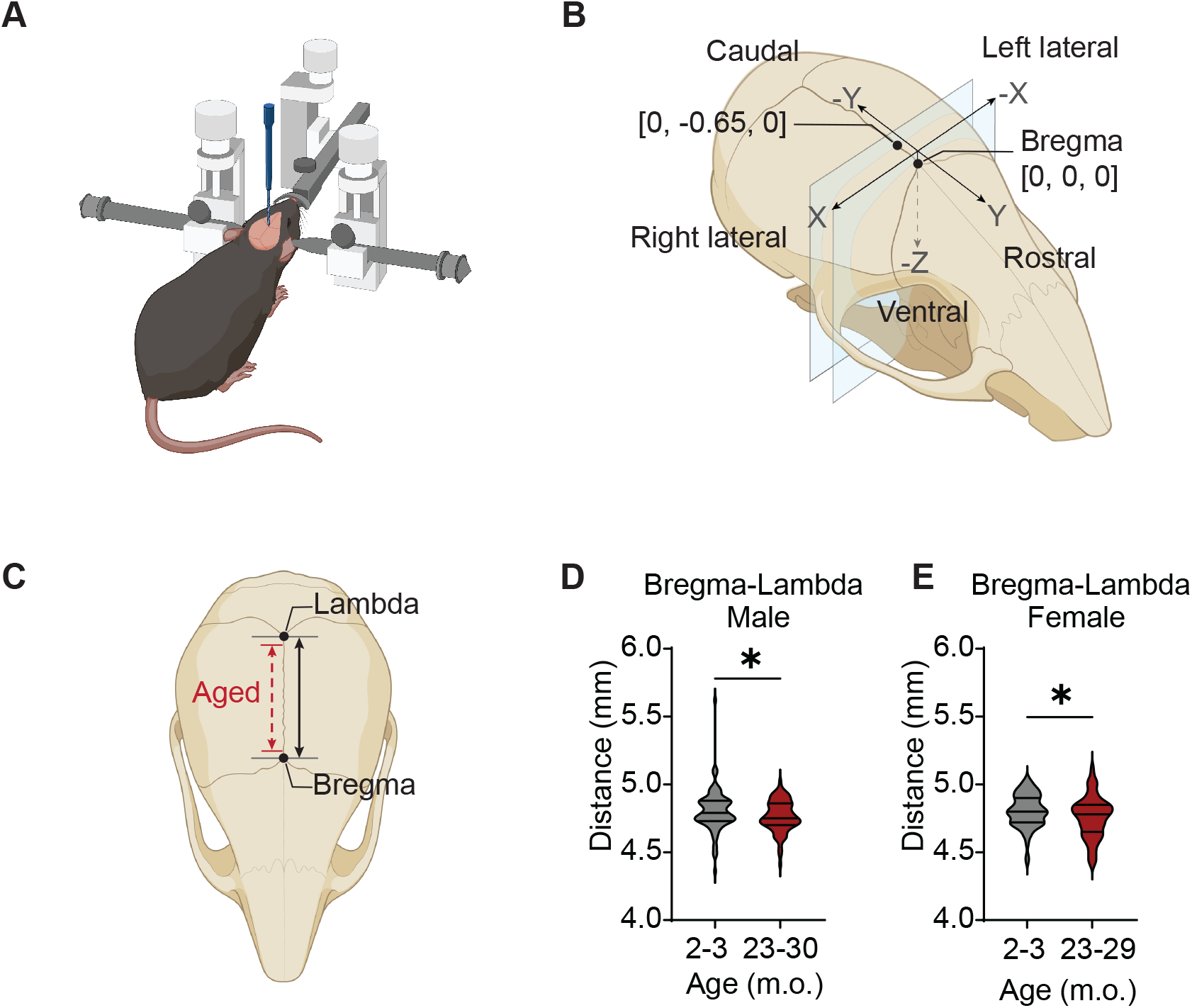
Age-related changes in bregma-lambda distance. (**A**) Stereotaxic setup used for acquisition of cranial measurements. (**B**) Oblique top-lateral view of the mouse skull showing orientation along the rostrocaudal (Y), mediolateral (X), and dorsoventral (Z) axes, with bregma as the stereotaxic origin and measurement planes corresponding to panels **1C**, Fig **2A** and **2D**. (**C**) Dorsal view illustrating the bregma-lambda distance (black) and age-related shortening (red) (schematic, not to scale). (**D–E**) Violin plots of bregma-lambda distance in young and aged male (**D**) and female (**E**) mice. Horizontal lines indicate median and interquartile range. *P* < 0.05 by Welch’s t-test.

### Statistical analysis

Descriptive statistics (mean, standard deviation [SD], standard error of the mean [SEM], and coefficient of variation [CV]) were computed for each group. Unpaired comparisons of means between young and aged groups used Welch’s *t*-test (one-tailed for bregma-lambda distance, two-tailed for Z coordinates), selected *a priori* to account for unequal group variances. Equality of variances was assessed by F-test. To compare the proportional (percent) change between young and aged groups across sexes or anatomical sites, log response ratios (LRR) were computed for each condition as LRR = log(mean_aged_/mean_young_), and differences in LRRs were tested using the delta method with a Wald *z*-test. Robustness of confidence intervals and *P*-values was verified by nonparametric bootstrap resampling (20,000 resamples). Statistical significance was set at α = 0.05. Analyses were performed by GraphPad Prism and custom-written Python scripts.

## RESULTS

### Age-related shortening of the rostro-caudal skull axis

Bregma-lambda distance was significantly reduced with aging in both sexes (**Figure 1C-E**; **Table 1**). In males, the mean distance decreased from 4.803 ± 0.155 mm (mean ± SD; *n* = 89) in young animals to 4.764 ± 0.115 mm in aged animals (*n* = 87), a reduction of 0.81% (Welch’s one-tailed *t*-test, *P* < 0.03). In females, the mean distance decreased from 4.805 ± 0.121 mm (*n* = 80) to 4.758 ± 0.152 mm (*n* = 67), a reduction of 0.98% (Welch’s one-tailed *P* < 0.02). The magnitude of shortening did not differ significantly between sexes (log response ratio comparison, 95% CI included unity). Inter-individual variability diverged by sex: the CV was significantly reduced in aged males (3.22% to 2.41%; F-test, *P* = 0.006), whereas females showed a non-significant trend toward increased CV (2.52% to 3.20%; *P* = 0.05).

**Table 1.** Bregma-lambda distance in young (2–3 months) and aged (23–30 months) male and female C57BL/6J mice.

|  | Male |  | Female |  |
| --- | --- | --- | --- | --- |
|  | Young<br>(2–3 m) | Aged<br>(23–30 m) | Young<br>(2–3 m) | Aged<br>(23–30 m) |
| n | 89 | 87 | 80 | 67 |
| Mean (mm) | 4.803 | 4.764 | 4.805 | 4.758 |
| SD (mm) | 0.155 | 0.115 | 0.121 | 0.152 |
| SEM (mm) | 0.016 | 0.012 | 0.014 | 0.019 |
| CV (%) | 3.22 | 2.41 | 2.52 | 3.20 |
| % change vs. young | - 0.81% |  | - 0.98% |  |
| Welch's t-test | P < 0.03 |  | P < 0.02 |  |
| F test (variances) | P = 0.006 |  | P = 0.05 (n.s.) |  |

*n*, number of animals; *CV*, coefficient of variation; *n.s.*, not significant. *SD*, standard deviation; *SEM*, standard error of the mean. Percent change is calculated relative to the young group mean. One-tailed Welch's *t*-test; *F* test for equality of variances.

### Age-related flattening of the rostral-lateral calvaria

The dorsoventral (Z) coordinate at the rostral-lateral site (X = ±2.5 mm, Y = 0) shifted toward less negative values with aging in both sexes (**Figure 2A-C**; **Table 2**). In males, Z values increased from -0.387 ± 0.058 mm (left) and -0.383 ± 0.055 mm (right) to -0.356 ± 0.050 mm and - 0.355 ± 0.050 mm, corresponding to relative reductions in dome depth of 7.99% and 7.18%, respectively (both *P* < 0.001). In females, Z values increased from -0.379 ± 0.055 mm (left) and - 0.376 ± 0.054 mm (right) to -0.346 ± 0.042 mm and -0.347 ± 0.041 mm, representing reductions of 8.51% (left) and 7.73% (right; both *P* < 0.001). The magnitude of flattening did not differ significantly between sexes at this site (*P* = 0.84, both sides). Variance was significantly reduced in aged females (F-test, *P* = 0.02, both sides) but not in aged males (*P* > 0.19).

**Table 2.** Dorsoventral coordinate (Z) at the rostral-lateral calvarial site (X = ±2.5 mm, Y = 0) in young and aged male and female C57BL/6J mice.

|  | Male |  |  |  | Female |  |  |  |
| --- | --- | --- | --- | --- | --- | --- | --- | --- |
| | Left ($X = -2.5$) | | Right ($X = +2.5$) | | Left ($X = -2.5$) | | Right ($X = +2.5$) | |
|  | Young | Aged | Young | Aged | Young | Aged | Young | Aged |
| n | 88 | 87 | 88 | 87 | 107 | 63 | 107 | 63 |
| Mean Z (mm) | -0.387 | -0.356 | -0.383 | -0.355 | -0.379 | -0.346 | -0.376 | -0.347 |
| SD (mm) | 0.058 | 0.050 | 0.055 | 0.050 | 0.055 | 0.042 | 0.054 | 0.041 |
| CV | 14.91 | 14.04 | 14.40 | 13.98 | 14.54 | 12.21 | 14.42 | 11.85 |
| % change | 7.99% |  | 7.18% |  | 8.51% |  | 7.73% |  |
| Welch's t-test | $P < 0.001$ | | $P < 0.001$ | | $P < 0.001$ | | $P < 0.001$ | |
| F test (var.) | $P = 0.19$ | | $P = 0.33$ | | $P = 0.02$ | | $P = 0.02$ | |

*n*, number of animals; CV, coefficient of variation; *n.s.*, not significant; SD, standard deviation.
Percent change is calculated relative to the young group mean. Two-tailed Welch's *t*-test; *F* test
for equality of variances.

**Figure 2.**
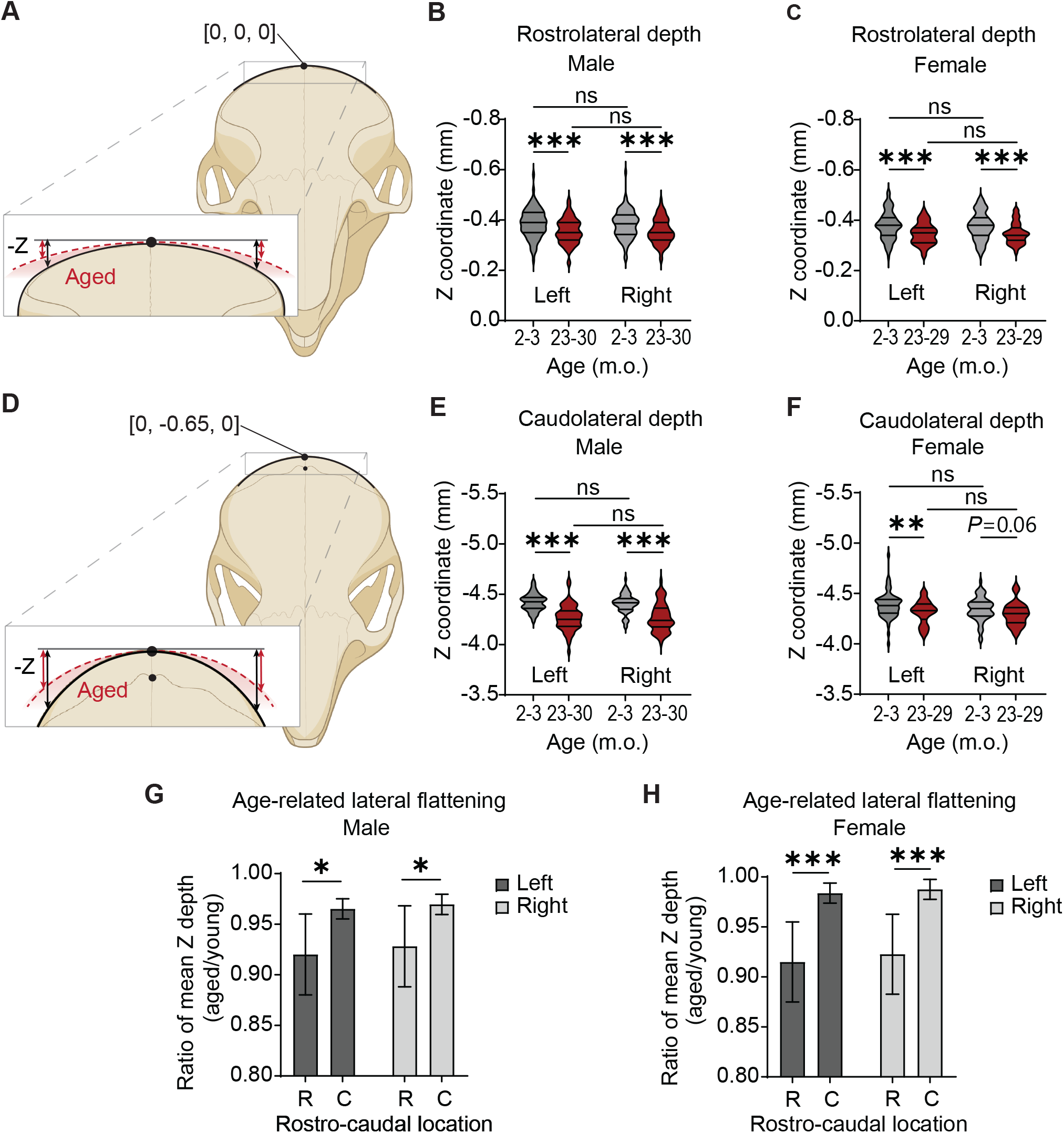
Age-related changes in lateral calvarial height in mice. (**A**) Frontal view with inset illustrating dorsoventral (Z) coordinate measurements at the rostral-lateral site X = ±2.5, Y = 0) in young (black) and aged mice (red) (schematic, not to scale). (**B-C**) Violin plots of Z coordinates at the rostral-lateral site in young and aged male (**B**) and female (**C**) mice. *P < 0.001 by Welch’s t-test; ns, not significant. (**D**) Frontal view with inset illustrating dorsoventral (Z) coordinate measurements at the caudal-lateral site X = ±2.5, Y = −0.65) in young (black) and aged mice (red) (schematic, not to scale). (**E-F**) Violin plots of Z coordinates at the caudal-lateral site in young and aged male (**E**) and female (**F**) mice. *P < 0.01, **P < 0.001 by Welch’s t-test; ns, not significant. (**G-H**) Bar graphs showing Z coordinates in aged mice relative to young mice at the rostral (R, Y = 0) and caudal (C, Y = −0.65) sites for males (**G**) and females (**H**). Left (X = −2.5) and right (X = +2.5) measurements are shown. Error bars denote 95% confidence intervals. *P < 0.05 and ***P < 0.001 by delta-method inference using log response ratios.

### Attenuated but significant flattening of the caudal-lateral calvaria, with sex differences

At the more caudal-lateral site (X = ±2.5 mm, Y = −0.65 mm), Z coordinates also shifted toward less negative values with aging, but to a significantly lesser extent than at the rostral site (**Figure 2D-F**; **Table 3**). In males, Z values increased from -4.419 ± 0.086 mm (left) and -4.406 ± 0.093 mm (right) to -4.265 ± 0.129 mm and -4.272 ± 0.139 mm, corresponding to reductions in depth of 3.48% (left) and 3.04% (right; both *P* < 0.001). In females, the left side shifted from -4.383 ± 0.131 mm to -4.315 ± 0.114 mm (1.62%; *P* = 0.009), while the right side changed from -4.342 ± 0.116 mm to - 4.296 ± 0.114 mm (1.24%; *P* = 0.06). Log response ratio analysis confirmed that caudal-lateral flattening was significantly greater in males than females (*P* < 0.01, both sides). In contrast to the rostral site, the variance at the caudal-lateral site increased significantly in aged males (F-test, *P* = 0.002, both sides) while remaining unchanged in females.

**Table 3.** Dorsoventral coordinate (Z) at the caudal-lateral calvarial site (X = ±2.5 mm, Y = −0.65 mm) in young and aged male and female C57BL/6J mice.

|  | Male |  |  |  | Female |  |  |  |
| --- | --- | --- | --- | --- | --- | --- | --- | --- |
| | Left (X = $-2.5$) | | Right (X = $+2.5$) | | Left (X = $-2.5$) | | Right (X = $+2.5$) | |
|  | Young | Aged | Young | Aged | Young | Aged | Young | Aged |
| n | 64 | 58 | 64 | 58 | 61 | 36 | 61 | 36 |
| Mean Z (mm) | -4.419 | -4.265 | -4.406 | -4.272 | -4.383 | -4.315 | -4.342 | -4.296 |
| SD (mm) | 0.086 | 0.129 | 0.093 | 0.139 | 0.131 | 0.114 | 0.116 | 0.114 |
| CV | 1.96 | 3.02 | 2.12 | 3.25 | 3.00 | 2.64 | 2.68 | 2.65 |
| % change | 3.48% |  | 3.04% |  | 1.62% |  | 1.24% |  |
| Welch's t-test | P < 0.001 |  | P < 0.001 |  | P = 0.009 |  | P = 0.06 |  |
| F test (var.) | P = 0.002 |  | P = 0.002 |  | P = 0.37 |  | P = 0.92 |  |

325 *n*, number of animals; CV, coefficient of variation; *n.s.*, not significant; SD, standard deviation. Percent change is calculated relative to the young group mean. Two-tailed Welch's *t*-test; *F* test for equality of variances.

## DISCUSSION

The present study provides a systematic, large-sample characterization of age-related change in surgically relevant calvarial geometry in male and female C57BL/6J mice. Three findings carry direct relevance for stereotaxic neurosurgery in aged rodents.

First, the bregma-lambda distance was reduced by ∼0.8-1.0% in both sexes, consistent with age-related shifts toward reduced calvarial appositional growth and net resorption across the sutures (Manolagas 2010). Age-related cortical atrophy and ventricular enlargement in mice (Taylor et al. 2020) may further reduce the intracranial expansile pressure that sustains calvarial outward growth (Pedersen et al. 2018; Robling et al. 2006). Consistent with altered skull geometry in aged mice, independent studies that involve stereotaxic targeting of hypothalamus and striatum empirically discovered that higher rostro-caudal (Y-axis) coordinates were required in older animals (Tokizane et al. 2024; Friedman et al. 2020). The bregma-lambda shortening quantified here provides a geometric basis for this empirical shift, as a shorter skull places anatomical brain structures at systematically different absolute coordinates, often higher, relative to bregma.

Second, the rostral-lateral calvaria (X = ± 2.5 mm, Y = 0), the reference site for skull-flat leveling, undergoes a larger age-related flattening in both sexes (∼7-9% reduction in dorsoventral depth) compared to the caudal-lateral site (X = ± 2.5 mm, Y = - 0.65 mm, ∼1-3%) (**Figure 2G-H**). This establishes a significant anteroposterior gradient of calvarial change, and may reflect regional differences in periosteal neurovascular architecture, which decline earlier in the frontal bone than in the parietal bone with aging in the mouse (Horenberg et al. 2025), or differential sensitivity to reduced intracranial pressure from brain atrophy across the calvaria. In addition, the resulting ∼0.03 mm bilateral reduction in dome height at the rostral-lateral calvaria introduces a systematic flatness at the leveling site; for deep or laterally displaced targets, even small angular offsets can accumulate into meaningful dorsoventral displacement errors (Rangarajan et al. 2016).

Aging was also associated with a sexually divergent reorganization of calvarial morphology and inter-individual variability. In males, variability in the bregma-lambda distance decreased with age, while caudal-lateral depth variability increased; in females, midline length variability trended upward and rostral-lateral depth variability decreased. Notably, age-related caudal-lateral flattening was significantly greater in males than in females, consistent with sex-specific remodeling trajectories in C57BL/6 mice. Prior studies report more abrupt age-related changes in estrogen-responsive skeletal remodeling programs in females than in males (Glatt et al. 2007), and estrogen withdrawal has been linked to increased heterogeneity in intramembranous bone remodeling, including the calvaria (Khosla et al. 2012; Manolagas 2010). How these sex-specific remodeling processes contribute to region-specific changes in calvarial geometry and variability with aging remains to be determined.

A consistent left > right asymmetry in Z-coordinate change likely reflects a systematic procedural bias such as preferred head positioning in the stereotaxic frame by a single operator, rather than true anatomical lateralization and therefore warrants independent verification. Study limitations include sampling only two lateral positions and the absence of more posterior measurements; whether the observed calvarial changes translate into quantifiable targeting errors remains to be tested directly. Nonetheless, these data provide a quantitative framework for coordinate adjustment and support routine pilot histological verification when working in aged cohorts.

Together, these findings demonstrate that aged mouse skull geometry departs measurably from the young-adult baseline on which standard atlases are based. Laboratories performing stereotaxic surgery in aged mice should empirically validate skull-flat leveling and adjust rostro-caudal coordinates for age-related skull shortening. These results highlight the need for age-specific cranial reference data in rodents.

## ACKNOWLEDGMENTS

We thank Lucy Culp for illustrations of the mouse skull and anatomical landmarks included in the figures, Charles Fisher for assistance with data organization, Michael Lyons for helpful discussions, and members of the Knight lab for technical assistance. Figure 1A was created with BioRender.com. This work was supported by the National Institutes of Health (R01-DK106399 and R01-DK138127 to Z.A.K; T32-AG000266 and F32-AG063488 to H.J.) and the Glenn Foundation for Medical Research (to H.J.). Zachary A. Knight is an Investigator of the Howard Hughes Medical Institute.

## AUTHOR CONTRIBUTIONS

H.J.: Conceptualization, methodology, data collection, formal analysis, writing-original draft, review and editing; Z.A.K.: Supervision, writing-review and editing.

## DATA AVAILABILITY STATEMENT

The data that support the findings of this study are available from the corresponding author upon reasonable request.

## ETHICS STATEMENT

All procedures were approved by the IACUC of UCSF (Protocol # AN179674-02A) and conducted in accordance with the Guide for the Care and Use of Laboratory Animals (8th ed.).

